# Spring phenology dominates over shade in affecting seedling performance and plant attack during the growing season

**DOI:** 10.1101/2020.01.05.895086

**Authors:** R. W. McClory, L. J. A. van Dijk, J. Mutz, J. Ehrlén, A. J. M. Tack

## Abstract

1. Climate change is affecting both the abiotic environment and the seasonal timing of life history events, with potentially major consequences for plant performance and plant-associated food webs. Despite this, we lack insights into how effects of plant phenology on plant performance and food webs depend on environmental conditions, and to what extent effects of phenology and the environment on plant performance are direct vs. mediated by changes in the plant-associated community.
2. We conducted a multifactorial field experiment to test for the effect of spring phenology and shade on *Quercus robur* seedling traits and performance, as well as attacks by specialist plant pathogens, insects and small mammals.
3. Spring phenology strongly affected seedling performance whereas shade only affected leaf thickness and chlorophyll. Likewise, spring phenology strongly affected herbivore and pathogen attack, whereas shade and its interaction with spring phenology only explained a minor part of the variation. Small mammals preferentially attacked later phenology seedlings, which strongly affected plant survival, while insect herbivores and pathogens did not mediate the effect of spring phenology and shade on plant performance.
4. ***Synthesis:*** This study highlights that the effect of spring phenology outweighs the effect of environmental context on plant performance and plant attack during the growing season. Interestingly, small mammal herbivores, and not diseases and insect herbivores, may play a key role in mediating the effect of spring phenology on plant performance. Together, these findings advance our understanding of the consequences of climate-induced changes in spring phenology and the abiotic environment on plant performance within a community context.

## Introduction

The abiotic environment plays a key role in creating spatial and temporal variation in spring phenology, with major consequences for organism performance, species interactions and food web structure (Kharouba et al., 2018; Thackeray et al., 2016). However, we lack studies that disentangle the joint effects of spring phenology and the abiotic environment during the remaining part of the growing season on organism performance and species interactions (Routhier & Lapointe, 2002). Such knowledge is important to predict the effects of climate change, as both the phenology of species and the abiotic context are expected to change. For example, early budburst may be disadvantageous in years with late frosts, whereas in other years early-phenology plants may benefit from a longer growing season (Bennie, Kubin, Wiltshire, Huntley, & Baxter, 2010). We also lack insights in the extent to which the effects of spring phenology and the abiotic environment influence plant performance directly, and to what extent these effects are mediated by the plant-associated food web.

Spring phenology and the abiotic environment during the growing season may interactively shape both plant performance and the presence and abundance of its associated species. For example, early spring phenology can increase or decrease oak growth, whereas shade decreases growth (Verdú & Traveset, 2005; Welander & Ottosson, 1998). Pathogen infection is well-known to be directly affected by abiotic environmental conditions such as temperature and humidity (Austin & Wilcox, 2012; Elad, Messika, Brand, David, & Sztejnberg, 2007), but also requires synchrony between susceptible plant tissue and spore release (Guest & Brown, 1997). Spring phenology has been shown to affect insect and mammal herbivory, with early bud-burst being associated with both lower (Barber & Fahey, 2015; Pearse, Funk, Kraft, & Koenig, 2015) and higher (Crawley & Akhteruzzaman, 1988; Heimonen et al., 2017) levels of herbivory. Despite evidence for independent and often contrasting effects of spring phenology and shade, we lack studies on how these factors interact to influence plant performance and attack.

Spring phenology and the abiotic environment may also affect plant performance indirectly via effects on the plant’s herbivores and pathogens. In such cases, changes in climate may directly affect plant attack, which indirectly lowers plant performance. Disentangling the direct and indirect nature of the interactive effect of spring phenology and the abiotic environment on plant performance seems particularly relevant, as a changing climate is expected to impact both spring phenology and the abiotic environment. Such assessments of the potential consequences for plant performance also need to account for that the abundance of different types of plant attackers are commonly negatively or positively correlated. For example, a recent meta-analysis showed that the presence of pathogens can strongly affect the preference and performance of herbivores (Fernandez-Conradi, Jactel, Robin, Tack, & Castagneyrol, 2018), while herbivores can affect the incidence and severity of plant pathogens in turn (Abdala-Roberts et al., 2019; Stout, Fidantsef, Duffey, & Bostock, 1999; Szczepaniec & Finke, 2019). However, the impact of such multi-attacker combinations for plant performance are unclear (Hauser, Christensen, Heimes, & Kiaer, 2013).

The overarching aim was to identify the impact of spring phenology and shade on the performance of seedlings of the pedunculate oak, *Quercus robur*, as well as on attack by its associated pathogens and herbivores. To achieve this, we conducted a multifactorial field experiment manipulating date of seedling germination and shade level, and measured plant traits, performance, and rates of attack by fungal pathogens, herbivorous insects and small mammals. More specifically, we addressed the following questions:

1. How do spring phenology and shade affect seedling performance?
2. How do spring phenology and shade affect the incidence and severity of plant attack?
3. Do attacks by pathogens and herbivores mediate the effect of phenology and shade on seedling performance?
4. Are rates of attack by insect herbivores and fungal pathogens correlated? In other words, are high rates of insect herbivory associated with high or low rates of pathogen infection?

We expected early spring phenology to increase, and shade to decrease, plant performance. Due to the high susceptibility of young oak leaves to infection by powdery mildew, we expected that late seedlings (which develop when pathogen spore loads are higher) would have higher levels of pathogen infection. We hypothesized that the independent and interactive effects of spring phenology and shade on plant performance take place through two non-mutually exclusive mechanisms: i) by directly affecting plant performance; and ii) by affecting plant attackers, which indirectly affect plant performance. Finally, because insect herbivores frequently avoid plants that have high pathogen infection levels (Fernandez-Conradi et al., 2018), we expected a negative correlation between severity of pathogen infection and insect herbivory.

## Materials and Methods

### Study system

The pedunculate oak, *Quercus robur*, is a deciduous tree species belonging to the family Fagaceae. The pedunculate oak is common throughout Europe and reaches the northern limit of its range in southern Norway, Finland and Sweden (Zanetto, Roussel, & Kremer, 1994). In Sweden, the pedunculate oak is a dominant tree species, and hosts numerous herbivorous insects (Ranius & Hedin, 2001) and pathogens (Horst, 2013). The phenology of the pedunculate oak varies strongly within and among populations and between years (Bacilieri, Ducousso, & Kremer, 1995; Kremer, Le Corre, Petit, & Ducousso, 2010).

In Europe, the most common fungal pathogens on the pedunculate oak are powdery mildews. In Sweden, the oak powdery mildew complex is dominated by two species belonging to the genus *Erysiphe.* The two species are spatially separated across the leaf surface: *E. alphitoides* mainly infects the upper leaf surface, and *E. hypophylla* is restricted to the lower leaf surface (Desprez-Loustau et al., 2018). Infection may cause tissue necrosis and is potentially devastating for young oak seedlings in natural systems and in tree nurseries (Marçais & Bréda, 2006). Infection in spring starts from overwintering sexual spores. During the growing season, the pathogen produces wind-dispersed, asexual spores (Marçais & Desprez-Loustau, 2014). The pedunculate oak also harbors a large number of herbivorous insects, including several species of Lepidoptera (Southwood, 1961). Further, oak seedlings are regularly browsed upon by various small mammals such as mice, voles and hares (Jensen, Götmark, & Löf, 2012).

### Experimental design

To identify how variation in spring phenology and shade can impact *Q. robur* seedling performance, as well as their associated pathogens and herbivores, we manipulated spring phenology and shade in a multifactorial design (Fig. S1). In order to create phenological differences between oak seedlings, acorns were planted at three-week intervals: i) early phenology acorns were planted on 22 April 2018; ii) medium phenology acorns were planted on 15 May 2018; and iii) late phenology acorns were planted on 3 June 2018. Acorns were planted in plastic pots (7 cm × 7 cm × 18 cm) with potting soil (Krukväxtjord, SW Horto, Hammenhög, Sweden) and placed in a greenhouse (21°C day/18°C night). Seedlings were translocated to the field directly after germination to ensure natural exposure to infection and herbivory during their entire growth period. Seedlings were kept within the pots to prevent the confounding effect of spatial variation in soil types, and pots were buried into the ground to prevent heat damage from sunlight. A ground sheeting (Fågelskrämma, Stockholm, Sweden) was placed around the pots to prevent competition with other plants. In the field, seedlings from each phenology treatment were divided equally into light shade (45% of light blocked) and heavy shade (65% of light blocked) treatment groups, with six blocks (5 m × 3 m) for each shade treatment. The field site was near the Bergius Botanical Garden (N 59°22’3.023”, E 18°3’3.907”). An electric fence (Gallagher, Stockholm, Sweden) was established around the field site to exclude large herbivores, such as deer. The seedlings were watered ad libitum throughout the experiment, taking care to keep soil moisture constant across treatment combinations.

### Data collection

To study how seedling traits and performance were influenced by spring phenology and shade, we measured several responses related to plant physiology and size. We measured leaf thickness (recorded with an IP-54 Electronic Outside Micrometre, Helios Pressier, Germany) and leaf chlorophyll (recorded as chlorophyll content index [CCI], with a CCm-200+ chlorophyll meter, Optisciences, Hudson, USA) on 8 August 2018, seedling height on 15 August 2018, leaf size on 20 August 2018 and the total number of leaves on 23 August 2018.

To study how spring phenology and shade affect attack by pathogens and herbivores, we measured powdery mildew infection on four occasions (25 June, 4 July, 17 July and 31 July 2018), and leaf herbivory on two occasions (25 June and 19 July 2018). For all seedlings, we recorded the presence or absence of powdery mildew and herbivore damage on each leaf (referred to as “infection incidence” and “herbivory incidence” respectively). We estimated powdery mildew severity as the percentage of each leaf on a seedling covered by the pathogen, with the upper and lower surfaces of the leaf measured separately. Likewise, we scored leaf herbivory severity as the percentage damage per leaf (Johnson, Bertrand, & Turcotte, 2016). Severity scores were then averaged among all of a seedling’s leaves to get an average severity of herbivore and pathogen attack per seedling.

To investigate the impact of spring phenology and shade on attack by small mammals, we recorded feeding marks made by small mammals. Small mammal feeding marks were easy to identify and were characterized by heavy feeding on the acorn and/or seedling, often with the consumption of the entire seedling.

### Statistical analyses

Statistical analyses were performed using R v 1.2.1335 (RStudio Team, 2018). Model structures, response variables and transformations are summarized in Table S1.

#### Effects of spring phenology and shade on seedling performance

To assess the impact of spring phenology and shade on seedling traits and performance, we used a linear mixed effect model, as implemented with the *lmer* function in the *lme4* package (Bates, Maechler, Bolker, & Walker, 2014). More specifically, we modelled leaf chlorophyll, leaf thickness, seedling height, leaf area, leaf number and survival as a function of the fixed effects *‘phenology’*, *‘shade’* and *‘phenology* × *shade’*. To account for environmental variation between blocks, the random factor *‘block’* was included in the model.

#### Effects of spring phenology and shade on plant attack

To assess the impact of phenology and shade on the leaf-level incidence of powdery mildew and herbivory, we used a repeated-measures generalized linear mixed effects model with a binomial distribution using the *glmer* function in the *lme4* package (Bates et al., 2014). To assess the impact of phenology and shade on the average severity of powdery mildew and leaf herbivory per seedling, we used a repeated measures linear mixed effects model using the *lmer* function in the *lme4* package (Bates et al., 2014). We modelled the incidence and severity of powdery mildew (separately for the lower and upper leaf surface) and herbivory as a function of the fixed effects *‘phenology’*, *‘shade’* and *‘date’*. To account for any changes in treatment effects through time, we included interactions between date and phenology and shade. For incidence models, we included the random effect *‘plant ID’*, as incidence of attack was recorded for multiple leaves from the same seedling. We further included the random factor *‘block’*.

As the treatment effects changed through time (i.e., there were significant *‘phenology* × *date’* and/or *‘shade* × *date’* interactions), we created date-specific models using the functions *glmer* and *lmer* in the *lme4* package (Bates et al., 2014). More specifically, we modelled the date-specific incidence and severity of infection and herbivory as a function of the fixed effects *‘phenology’*, *‘shade’*, the interaction *‘shade* × *phenology’* and the random factors *‘block’* and *‘plant ID’*.

#### Effects of phenology and shade on seedling performance as mediated by plant attack

To test whether the effects of spring phenology and shade were mediated by plant attack, we compared models with vs. without the covariates powdery mildew infection and herbivory. Powdery mildew damage on the upper and lower leaf surface was expressed as the area under the disease progression curve (AUDPC), which gives a quantitative summary of disease intensity over time (Madden, Hughes, & van den Bosch, 2017). For herbivory, we averaged the percentage of leaf herbivory across the two recording dates. We modelled the seedling performance traits leaf chlorophyll, leaf thickness, seedling height, leaf area, leaf number and survival as a function of the fixed effects *‘phenology’*, *‘shade’*, *‘shade* × *phenology’*, *‘upper leaf AUDPC’, ‘lower leaf AUDPC’* and *‘herbivory’*. To account for environmental variation between blocks, the random factor *‘block’* was included in the model. Differences in the estimated effects of phenology and shade between the models with and without the covariates (i.e. powdery mildew infection and herbivory) would provide support for the hypothesis that the effects of spring phenology and shade on seedling physiology and growth are mediated by powdery mildew infection and/or leaf herbivory.

#### Correlation among different types of plant attack

To examine the relationship between different types of plant attack, we used a correlation analysis. We first calculated Kendall’s rank correlation coefficient between the AUDPC of powdery mildew on the lower leaf surface, AUDPC of powdery mildew on the upper leaf surface, and the averaged damage by insect herbivores using the observational data. As interactions between the types of attack can be obscured by different responses of attackers to spring phenology and shade, we also investigated the correlation between the powdery mildew and herbivory after accounting for spring phenology and shade. More specifically, we correlated the residuals from models where the upper and lower AUDPC and average herbivory damage were individually modelled as a function of the fixed effects *‘phenology’*, *‘shade’*, the interaction *‘shade* × *phenology’*, and the random factor *‘block’*.

## Results

### Effects of spring phenology and shade on seedling performance

Phenology affected seedling performance and leaf traits, with the exception of leaf thickness (Table 1). Seedlings with medium and late phenology had higher levels of chlorophyll and were taller (Fig. 1AB). Leaf area increased between early to late phenology seedlings, and medium phenology seedlings had the most leaves (Fig. 1CD). Shade only affected chlorophyll and leaf thickness (Table 1), with seedlings under light shade having lower chlorophyll levels and greater leaf thickness than seedlings under heavy shade (Fig. 1EF). The effect of spring phenology on plant performance and leaf traits did not differ among shade levels (i.e., there were no significant ‘*spring phenology* × *shade*’ interactions; Table 1). Seedling mortality strongly increased between early to late phenology groups, with 1.1% mortality in the early phenology group, 9.8% mortality in the medium phenology group, and 25.1% mortality in the late phenology group (Table 1).

**Figure 1.**
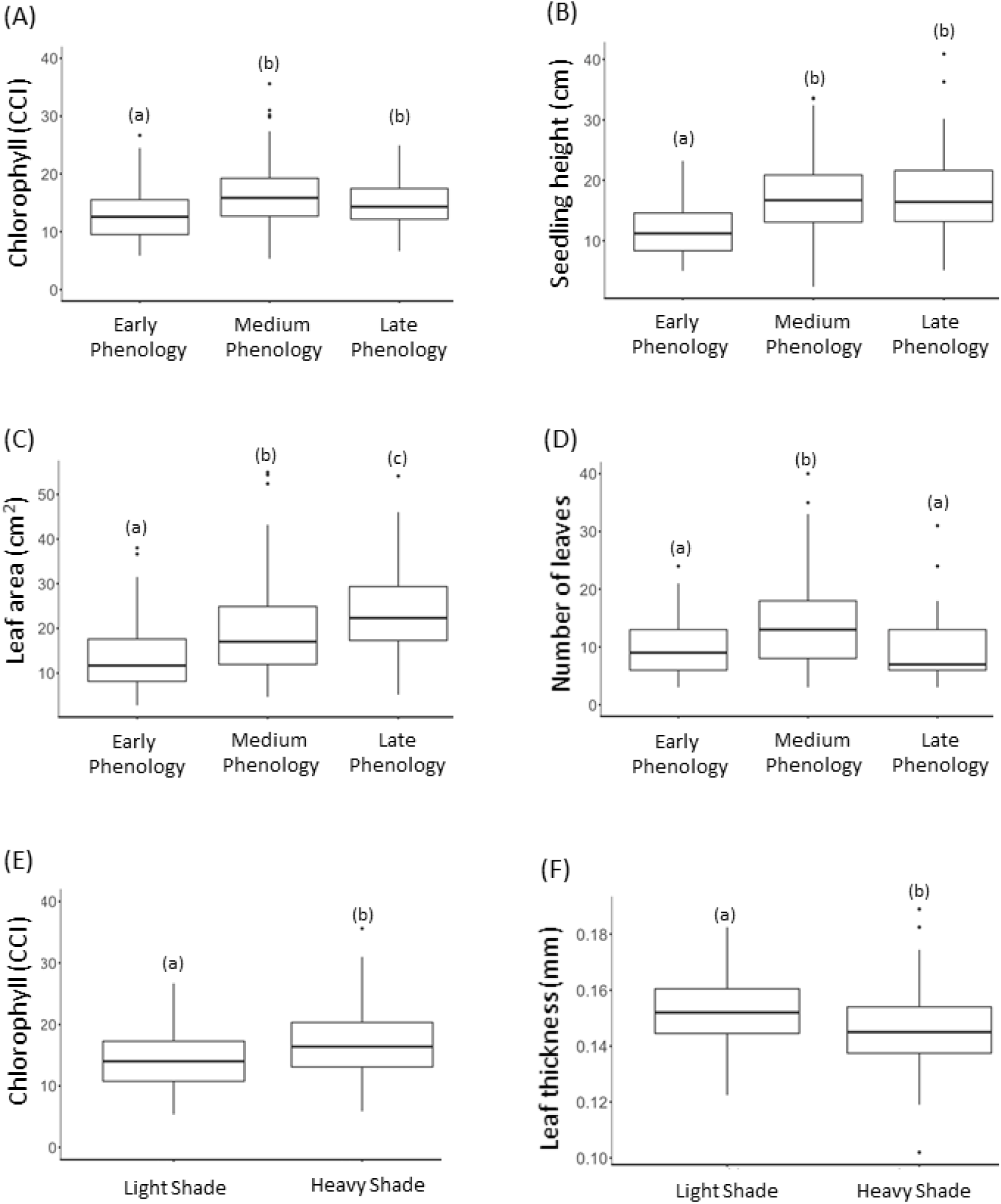
The impact of spring phenology and shade on performance traits of *Quercus robur* seedlings. Shown are the impact of spring phenology on (A) chlorophyll (recorded as chlorophyll content index [CCI]), (B) seedling height (C) leaf area and (D) number of leaves, and the impact of shade on (E) chlorophyll and (F) leaf thickness. The lowercase letters identify which groups are significantly different from each other (p<0.05) as based on post-hoc pairwise comparisons.

**Table 1.** The impact of phenology, shade and the interaction shade × phenology on performance and survival of oak seedlings (*Quercus robur*).

| Response variable | Phenology |  |  | Shade |  |  | Shade ×<br>Phenology |  |  |
| --- | --- | --- | --- | --- | --- | --- | --- | --- | --- |
| | Df | F/$\chi^2$ | P | Df | F/$\chi^2$ | P | Df | F/$\chi^2$ | P |
| Leaf thickness | 2 | 1.21 | 0.55 | 1 | 5.22 | **0.022** | 2 | 0.69 | 0.72 |
| Chlorophyll | 2 | 33.33 | **<0.0001** | 1 | 7.4 | **0.006** | 2 | 3.6 | 0.16 |
| Seedling height | 2 | 56.52 | **<0.0001** | 1 | 0.56 | 0.45 | 2 | 2.13 | 0.34 |
| Leaf area | 2 | 54.27 | **<0.0001** | 1 | 0.78 | 0.38 | 2 | 0.17 | 0.92 |
| Leaf number | 2 | 39.70 | **<0.0001** | 1 | 0.03 | 0.87 | 2 | 0.81 | 0.67 |
| Survival | 2 | 35.19 | **<0.0001** | 1 | 0.55 | 0.46 | 2 | 0.08 | 0.96 |

### Effects of spring phenology and shade on plant attack

Plant attack by small mammals was higher for the later-phenology seedlings: small mammals attacked 0% of the early phenology seedlings, 8.7% of the middle phenology seedlings, and 24.4% of the late phenology seedlings. Plant attack by small mammals was not affected by shade or its interaction with spring phenology (Table 2).

**Table 2.** The impact of phenology, shade and their interaction on small mammal attack, the proportion of a seedling’s leaves infected with powdery mildew on the upper leaf surface, the proportion of a seedling’s leaves infected with powdery mildew on the lower surface, and leaf herbivory, on *Quercus robur* seedlings

| Response variable | Week Number | Phenology |  |  | Shade |  |  | Shade × Phenology |  |  |
| --- | --- | --- | --- | --- | --- | --- | --- | --- | --- | --- |
| | | DF | F/$\chi^2$ | P | DF | F/$\chi^2$ | P | DF | F/$\chi^2$ | P |
| Small mammal attack | N/A | 2 | 32.8 | **<0.0001** | 1 | 1.07 | 0.59 | 2 | 0.2 | 0.9 |
| Upper leaf surface powdery mildew incidence | Week 1 | 2 | 2.49 | 0.29 | 1 | 0.13 | 0.72 | 2 | 0.93 | 0.63 |
|  | Week 2 | 2 | 41.85 | **<0.0001** | 1 | 0.11 | 0.74 | 2 | 2.5 | 0.29 |
|  | Week 4 | 2 | 3.4 | 0.18 | 1 | 4.87 | **0.027** | 2 | 1.58 | 0.45 |
|  | Week 6 | 2 | 7.05 | **0.029** | 1 | 2.4 | 0.12 | 2 | 1.84 | 0.4 |
| Lower leaf surface powdery mildew incidence | Week 4 | 2 | 67.12 | **<0.0001** | 1 | 0.29 | 0.59 | 2 | 0.58 | 0.75 |
|  | Week 6 | 2 | 60.15 | **<0.0001** | 1 | 0.16 | 0.69 | 2 | 2.45 | 0.29 |
| Herbivory incidence | Week 1 | 2 | 36.87 | **<0.0001** | 1 | 0.9 | 0.34 | 2 | 7.35 | **0.025** |
|  | Week 4 | 2 | 6.08 | **0.048** | 1 | 1.03 | 0.31 | 2 | 3.74 | 0.15 |

Spring phenology affected powdery mildew infection incidence on the upper and lower surface of the leaves, though the effect varied through time (Table S4). Directionality of this effect was inconsistent through time for the powdery mildew on the upper leaf surface (Table 2): in week 2, late phenology seedlings had the lowest infection incidence (Fig. 2A), whereas by week 6 infection incidence was lowest for the early phenology seedlings (Fig. 2A). On the lower leaf surface, infection incidence was consistently higher for later phenology seedlings (Table 2). Infection severity on the lower leaf surface, but not the upper leaf surface, was affected by phenology (Table S3; Fig. 2BD). In week 4, infection severity was higher for early phenology seedlings, but this pattern was no longer present by week 6 (Table S5; Fig 2D.). Towards the end of the experiment, infection incidence and severity were higher under heavy shade (Tables 2, S3 and S4; Fig.2EF). The effect of spring phenology on powdery mildew incidence and severity was not affected by the level of shade (i.e., the ‘*spring phenology* × *shade*’ interactions were not significant; Tables 2, S5).

**Figure 2.**
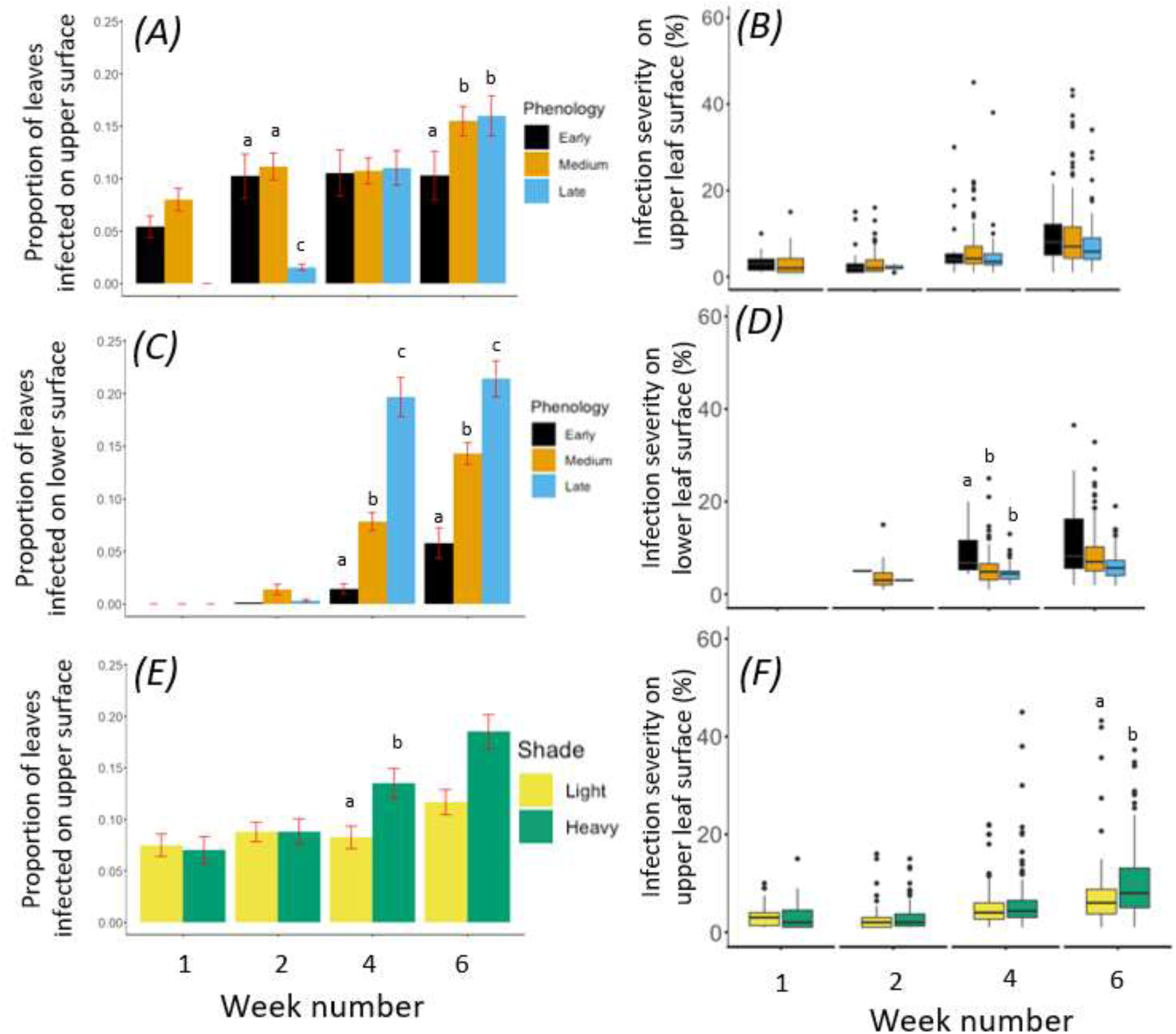
The impact of early, medium and late spring phenology and light and heavy shade treatments on *Quercus robur* seedlings. The impact of spring phenology on the proportion of leaves infected and severity of powdery mildew on (AB) the upper surface and (CD) lower surface of the leaf. The impact of shade on (E) the proportion of leaves infected by and (F) severity of powdery mildew on the upper surface of the leaf under light and heavy shade. The letters above the treatment levels identify which groups are significantly different from each other (p<0.05) as based on post-hoc pairwise comparisons.

Phenology affected herbivory incidence, but the pattern varied through time: herbivory was highest on medium-phenology seedlings in week 1, and highest for late-phenology seedlings in week 4 (Table 2; Fig. 3A). Moreover, the effect of spring phenology differed between the two shade treatments during week 1 (Table 2). Late phenology seedlings had greatly reduced herbivory incidence under heavy shade, compared to the same phenology group under light shade (Fig. 3B). There were no effects of spring phenology or shade on herbivory severity (Table S5).

**Figure 3.**
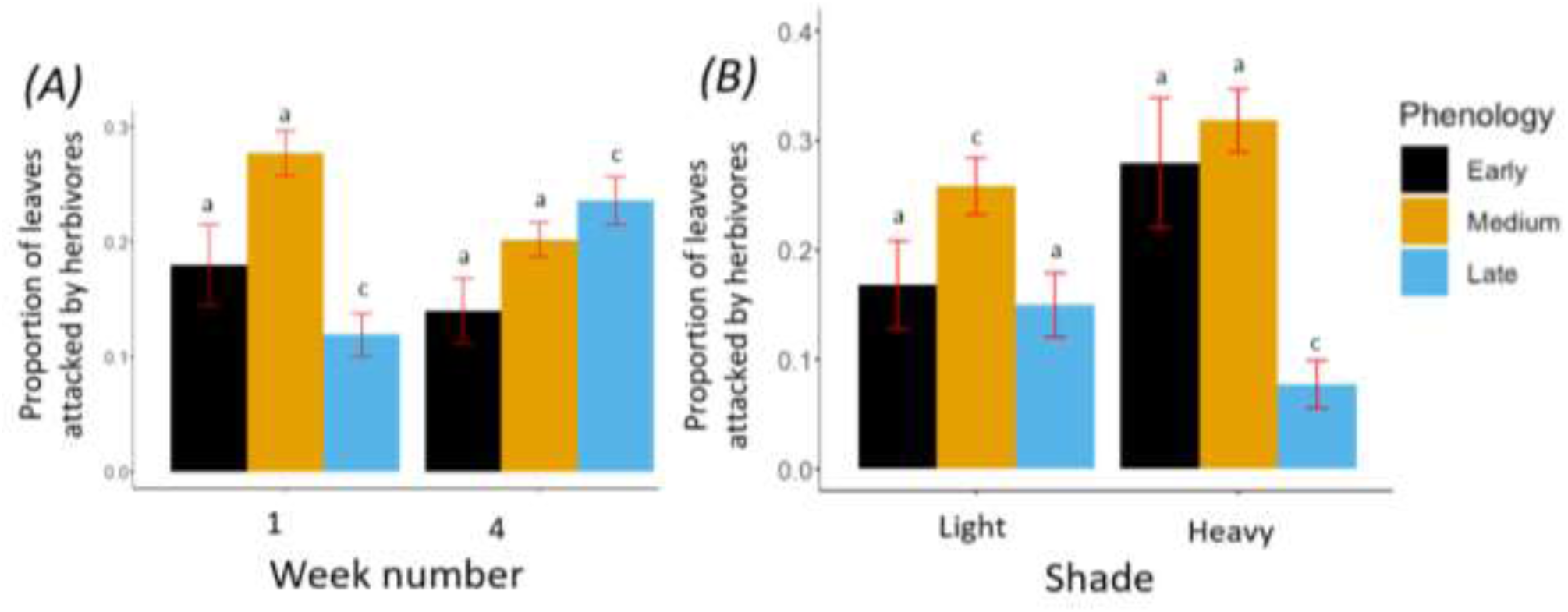
The impact of spring phenology and shade on the leaf herbivory incidence of *Quercus robur* seedlings. Panel A shows the impact of spring phenology on the proportion of leaves attacked by herbivores in week 1 and 4. Panel B shows the interaction of spring phenology and shade on the proportion of leaves attacked for the first week of the experiment. The lowercase letters identify which groups are significantly different from each other (p<0.05) as based on post-hoc pairwise comparisons.

### Effects of phenology and shade on seedling performance as mediated by plant attack

Small mammal herbivory was the major cause of seedling mortality: out of the 116 seedlings that died during the experiment, 109 seedlings died due to herbivory by small mammals. From those seedlings that were attacked by small mammals, only two individuals survived.

Powdery mildew and leaf herbivory did not mediate the effects of phenology and shade on seedling performance, as indicated by the lack of change in significance, and very minor change in effect estimates, of the terms *‘phenology’* or *‘shade’* when adding the covariates *‘upper leaf AUDPC’, ‘lower leaf AUDPC’* and *‘herbivory’* to the model (cf. Table 1 and Table S2). Leaf herbivory was only weakly positively related with leaf thickness, whereas powdery mildew infection was not associated with any of the seedling performance traits (Table S2).

### Correlation among different types of plant attack

No simple correlations were found between different types of plant attack. However, after accounting for differences in spring phenology and shade, we found a weak positive correlation between the upper leaf surface AUDPC score and average leaf herbivory (r_τ_ = 0.084, z = 2.52, df = 403, P = 0.012), indicating that disease and herbivory co-occurred more often than expected by chance. There was no relationship between AUDPC on the lower surface and leaf herbivory. There was a positive correlation between upper leaf surface AUDPC scores and lower leaf surface AUDPC scores (r_τ_ = 0.15, z = 4.48, df = 403, P <0.0001).

## Discussion

While several previous studies have focused on the impact of environmental variation in creating spatial variation in germination and bud burst, we are among the first to quantify the impact of spring phenology and environmental variation on plant performance and plant attack during the remainder of the growing season. We found that spring phenology had a major impact on seedling performance, whereas shade only affected leaf thickness and chlorophyll. Likewise, spring phenology had a major impact on plant attack, while the effects of shade were minor. While small mammals had a major effect on plant survival by preferentially attacking later phenology seedlings, insect herbivores and pathogens did not mediate the effect of spring phenology and shade on plant performance. The different types of plant attack were only weakly correlated. Taken together with the results of previous studies, our findings indicate that while abiotic factors like shade can have a major impact on spring phenology and leaf traits, spring phenology plays a more important role in shaping seedling growth and plant attack during the remainder of the growing season.

### Effects of spring phenology and shade on seedling performance

We found that seedlings were larger in the medium and late phenology treatment, but that mortality was lowest for early phenology seedlings. Our finding of reduced growth of early phenology seedlings contrasts with previous studies that showed that earlier spring phenology enhances seedling growth and survival (Seiwa, 1997, 1998; Seiwa & Kikuzawa, 1996; Verdú & Traveset, 2005). One explanation for this difference may be the experimental exclusion of plant-plant competition in our experiment: under natural conditions, early phenology seedlings may gain a competitive advantage by growing before canopy closure and escape competition from other plants. Our experimental seedlings had shade nets and ground sheeting, which removed the potential for this competitive advantage (DePamphilis & Neufeld, 1989; Miller, Winn, & Schemske, 1994; Seiwa, 2000). As expected, seedlings under heavy shade produced thinner leaves with higher levels of chlorophyll, most likely as a strategy to more efficiently capture the sparser light under the heavy shade treatment (Jackson, 1967; Valladares, Martinez-Ferri, Balaguer, Perez-Corona, & Manrique, 2000).

### Effects of spring phenology and shade on plant attack

Spring phenology had a major impact on seedling attack by small mammalian herbivores, pathogen infection and herbivorous insects, whereas shade and its interaction with spring phenology only explained a minor part of the variation. The apparent preference of small mammals for later-phenology seedlings may be explained by temporal synchrony between acorn condition and high attack rates by small mammal herbivores: by the time the attacks occurred, the early phenology acorns were already starting to shrivel due to extraction of resources from the acorn by the seedling.

Late phenology seedlings had the lowest disease levels on the upper leaf surface during the early part of the season, whereas infection levels on both the lower and upper surface were highest on the later-phenology seedlings towards the end of the experiment. The difference in observed disease levels early in the season may be due to a time lag between leaf colonization and the appearance of the symptoms or differences in our ability to detect infection (e.g., early infections on the young developing leaves of late-phenology seedlings may have gone undetected). The findings of higher disease levels on the later-phenology seedlings at the end of the experiment matches our a priori prediction that plants developing their leaves when the pathogen spore load in the air is already high would experience higher infection levels. Seedlings growing under heavy shade experienced higher infection incidence and severity on the upper leaf surface. This disagrees with previous studies showing that high light levels increased infection levels of multiple powdery mildew species (Kelly, 2002; Newsham, Oxborough, White, Greenslade, & McLeod, 2000), but matches field observations on higher infection levels on oaks in forest habitats when compared with unshaded open fields (Ekholm, Roslin, Pulkkinen, & Tack, 2017). Alternatively, shade-induced changes in leaf structure (i.e. thinner leaves) may have resulted in higher susceptibility of infection from powdery mildew (Giertych & Suszka, 2010). Importantly, this finding suggests that shade may more strongly affect pathogens on the upper leaf surface, which makes sense given the fact that pathogens on the lower surface are already buffered from high surface temperatures, low relative humidity and UV radiation (Aust & Huene, 1986; Hewitt, 1974; Benoit Marçais & Desprez-Loustau, 2014).

Herbivory incidence was highest in early and mid-phenology seedlings at the start of the experiment, especially for those under heavy shade. This pattern had changed towards the end of the experiment in late July, when herbivory incidence was highest on later phenology seedlings and was similar among shade levels. Hence, while previous studies have found that earlier budburst results in increased levels of herbivory of adult oak trees at the end of the season (Pearse, Baty, Herrmann, Sage, & Koenig, 2015; Pearse, Funk, et al., 2015), we find that early phenology seedlings have lower – not higher – levels of cumulative herbivory at the end of the season. In contrast to earlier studies (Muth, Kluger, Levy, Edwards, & Niesenbaum, 2008), we did not find an independent effect of shade on herbivory. Future studies may explore the mechanistic underpinning of the interactive effect of shade and phenology in determining herbivory of seedlings. For example, seasonality, plant ontogeny, and shade level may each affect a seedling’s defenses and nutritional profile, and therefore its quality as a resource to insect herbivores.

### Effects of phenology and shade on seedling performance as mediated by plant attack

Small mammals mediated the effect of spring phenology on plant survival: early phenology seedlings largely escaped attack, middle phenology seedlings had average rates of attack, and nearly one out of four late phenology seedlings died due to feeding by small mammals. Matching these findings, Seiwa (1998) previously demonstrated that seedling attack by small mammals was lower for early phenology seedlings. As nearly every plant attacked by small mammals died, small mammals did not affect other aspects of seedling performance (e.g., growth).

In contrast to the strong effect of small mammal herbivory on seedling survival, pathogen infection and insect herbivory had negligible effects on seedling survival and performance, and thus did not mediate the effects of phenology and shade. The limited effect of pathogens and leaf herbivory on plant performance contrasts with other studies on both seedlings (Norghauer & Newbery, 2014; Solé et al., 2019) and adult plants (Marçais & Bréda, 2006; Pearse, Funk, et al., 2015). One explanation for the lack of an effect of pathogens and insects on seedling performance may be the availability of stored seed reserves, which could offset any immediate costs of infection and herbivory (Grime & Jeffrey, 1965). Although drawing upon stored reserves can affect future survival and performance (Pearse, Funk, et al., 2015; Sala, Hopping, McIntire, Delzon, & Crone, 2012), we found no effect of infection or herbivory on overwinter survival of our experimental seedlings (all P > 0.05). Taken together, while small mammals directly kill seedlings and thereby play a clear and direct role in natural selection for seedling phenology, the seedling may be able to compensate the effect of pathogen and insect attack on plant performance by tapping into the stored resources in the acorn, at least in the short term.

### Correlation among different types of plant attack

Strong positive or negative interactions among plant attackers may strongly influence the net effect of phenology and abiotic environment on plant performance. However, we only found a weak, positive correlation between powdery mildew infection on the upper leaf surface and damage by free-feeding herbivores. The finding of a positive correlation between these two types of plant attack matches the expectation of a trade-off between the jasmonate defence pathway, which is elicited by chewing herbivores, and the salicylate defence-related pathway, which is elicited by biotrophic pathogens (Thaler, Humphrey, & Whiteman, 2012). However, studies have found mixed support for this hypothesis (Fernandez-Conradi et al., 2018; Moreira, Abdala-Roberts, & Castagneyrol, 2018; Tack & Dicke, 2013), and the lack of a relationship between herbivory and infection on the lower leaf surface indicates that – if a trade-off is present – it acts differently on two closely related, obligate plant pathogens sharing the same leaf. The positive relationship between infection at the lower and upper leaf surface may either be due to shared preferences for plant traits, or due to plant mediated interactions between *E. alphitoides* and *E. hypophylla*. Experimental competition studies may be a promising way to shed light on the competitive and/or facilitative interaction between these two cryptic pathogen species sharing the same leaf.

### Conclusions

With a changing climate, it will be crucial to understand how climate-induced changes in spring phenology and the abiotic environment interactively influence plant performance and plant-associated food webs. Our study is among the first to investigate the simultaneous effects of spring phenology and shade on plant performance and attack during the growing season. We found that the effect of spring phenology outweighs the effect of the abiotic environment (i.e., shade) on seedling performance and several types of plant attack. Interestingly, our results suggest that small mammals, but not herbivorous insects and pathogens, mediate the effects of spring phenology on plant performance. Future studies may aim to generalize the strong effect of spring phenology on small mammal attack in other plant species and explore the underlying mechanism and consistency of small mammal preference. Interestingly, despite the previously demonstrated importance of shade for creating spatial variation in spring phenology, we found that seedling performance and attack were largely unaffected by shade. Hence, we conclude that abiotic factors like shade may contribute to variation in phenology early in the season, but that spring phenology – and not shade – goes on to influence plant performance and attack during the growing season.

## Acknowledgements

We thank Eleanor Child for help with setting up the experiment and recording data, and the Bergius Botanical Gardens and the Swedish Museum of Natural History for permission to use the land. This work was supported by funding from the Swedish Research Council (2015-03993) to AJMT.

## Author contributions

RWM, AJMT, JE and LJAvD conceived and designed the experiment. RWM conducted the empirical work. RWM analyzed the data. RWM wrote the first draft, and all authors contributed to the final manuscript.

## Data accessibility

Data associated with this manuscript will be archived in the Dryad Digital Repository upon acceptance.

