## Supplemental Tables and Figures for "Spring phenology dominates over shade in affecting seedling performance and plant attack during the growing season"

**Supporting information.** R. W. McClory, L. J. A. van Dijk, J. Mutz, J. Ehrlén and A. J. M. Tack.

Supplementary figures and tables presented in order of appearance within the manuscript

**Figure S1.** Schematic overview of the multifactorial design of the experiment, including the replicate numbers of *Quercus robur* seedlings for each phenology and shade level.

**Table S1.** A summary of the models fitted for analyses. For each model, we specify the response examined the fixed [F] and random [R] effects included, and the link function applied. For identity links, we used a Gaussian distribution, and for logit links, we used a binomial distribution.

**Table S2.** The impact of phenology, shade, the interaction shade × phenology, the area under the disease progression curve (AUDPC) and leaf herbivory on the physiology, growth and survival of oak seedlings (*Quercus robur*).

**Table S3.** Repeated measures models of the proportion of leaves infected with powdery mildew and the severity of the infection on the upper and lower surface of *Quercus robur* seedlings leaves, as well as leaf herbivory, as a function of phenology, shade and date, and the interactions between phenology × shade, date × phenology, date × shade and date × phenology × shade. From weekly measurements during the period from 26 June 2018 to 31 July 2018.

**Table S4.** The impact of phenology, shade and their interaction on the severity of powdery mildew infection on the upper surface of the leaf, severity of powdery mildew infection on the lower surface of the leaf and leaf herbivory severity, on *Quercus robur* seedlings

**Figure S1.** Schematic overview of the multifactorial design of the experiment, including the replicate numbers of *Quercus robur* seedlings for each phenology and shade level.

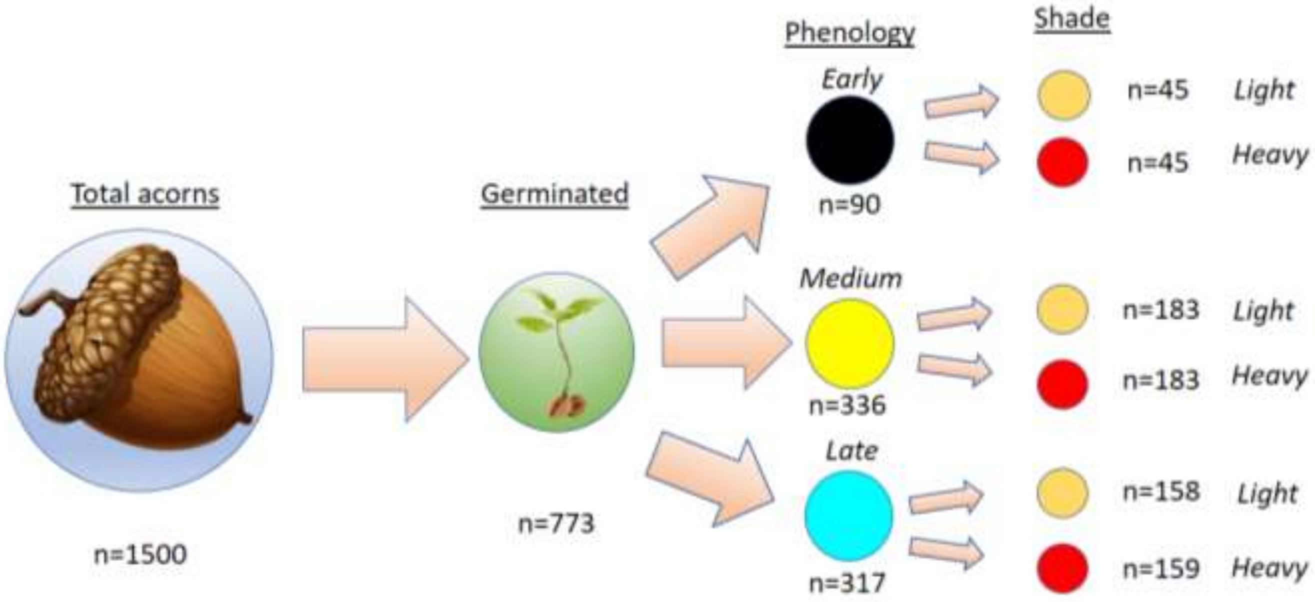

**Table S2.** A summary of the models fitted for analyses. For each model, we specify the response examined, the fixed [F] and random [R] effects included, and the link function applied. For identity links, we used a Gaussian distribution, and for logit links, we used a binomial distribution.

| Questions targeted | Response examined | Models fitted | Link |
|---|---|---|---|
| How do spring phenology and shade affect seedling performance and survival? | *Leaf thickness* | Phenology [F] + Shade [F] + Phenology × Shade [F] + Block [R] | Identity |
| | *Chlorophyll content*[1] | Phenology [F] + Shade [F] + Phenology × Shade [F] + Block [R] | Identity |
| | *Seedling height* | Phenology [F] + Shade [F] + Phenology × Shade [F] + Block [R] | Identity |
| | *Leaf area* | Phenology [F] + Shade [F] + Phenology × Shade [F] + Block [R] | Identity |
| | *Leaf number* | Phenology [F] + Shade [F] + Phenology × Shade [F] + Block [R] | Identity |
| | *Survival* | Phenology [F] + Shade [F] + Phenology × Shade [F] + Block [R] | Binomial |

| Questions targeted | Response examined | Models fitted | Link |
| --- | --- | --- | --- |
| How do spring phenology and shade affect plant attackers? | *Small mammal attack* | Phenology [F] + Shade [F] + Phenology × Shade [F] + Block [R] | Logit |
| | *Upper leaf surface powdery mildew infection incidence* (0/1) | i) Phenology [F] + Shade [F] + Phenology × Shade [F] + Date [F] + Phenology × Date [F] + Shade × Date [F] + Phenology × Shade × Date [F] + Block [R] + Plant_ID [R]<br>ii) Phenology [F] + Shade [F] + Phenology × Shade [F] + Block [R] + Plant_ID [R] | Logit |
| | *Lower leaf surface powdery mildew infection incidence* (0/1) | i) Phenology [F] + Shade [F] + Phenology × Shade [F] + Date [F] + Phenology × Date [F] + Shade × Date [F] + Phenology × Shade × Date [F] + Block [R] + Plant_ID [R]<br>ii) Phenology [F] + Shade [F] + Phenology × Shade [F] + Block [R] + Plant_ID [R] | Logit |
| | *Leaf herbivory incidence* (0/1) | i) Phenology [F] + Shade [F] + Phenology × Shade [F] + Date [F] + Phenology × Date [F] + Shade × Date [F] + Phenology × Shade × Date [F] + Block [R] + Plant_ID [R]<br>ii) Phenology [F] + Shade [F] + Phenology × Shade [F] + Block [R] + Plant_ID [R] | Logit |
| | *Upper leaf surface powdery mildew infection severity*[1] | i) Phenology [F] + Shade [F] + Phenology × Shade [F] + Date [F] + Phenology × Date [F] + Shade × Date [F] + Phenology × Shade × Date [F] + Block [R]<br>ii) Phenology [F] + Shade [F] + Phenology × Shade [F] + Block [R] | Identity |
| | *Lower leaf surface powdery mildew infection severity*[1] | i) Phenology [F] + Shade [F] + Phenology × Shade [F] + Date [F] + Phenology × Date [F] + Shade × Date [F] + Phenology × Shade × Date [F] + Block [R]<br>ii) Phenology [F] + Shade [F] + Phenology × Shade [F] + Block [R] | Identity |
| | *Leaf herbivory severity*[1] | i) Phenology [F] + Shade [F] + Phenology × Shade [F] + Date [F] + Phenology × Date [F] + Shade × Date [F] + Phenology × Shade × Date [F] + Block [R]<br>ii) Phenology [F] + Shade [F] + Phenology × Shade [F] + Block [R] | Identity |

| Questions targeted | Response examined | Models fitted | Link |
|---|---|---|---|
| Do powdery mildew and herbivory mediate the effect of phenology and shade on seedling performance and survival? | *Leaf thickness* | ii) Phenology [F] + Shade [F] + Phenology × Shade [F] + Upper AUDPC[F] + Lower AUDPC[F] + Herbivory[F] + Block [R] | Logit |
| | *Chlorophyll content*[1] | ii) Phenology [F] + Shade [F] + Phenology × Shade [F] + Upper AUDPC[F] + Lower AUDPC[F] + Herbivory[F] + Block [R] | Logit |
| | *Seedling height* | ii) Phenology [F] + Shade [F] + Phenology × Shade [F] + Upper AUDPC[F] + Lower AUDPC[F] + Herbivory[F] + Block [R] | Logit |
| | *Leaf area* | ii) Phenology [F] + Shade [F] + Phenology × Shade [F] + Upper AUDPC[F] + Lower AUDPC[F] + Herbivory[F] + Block [R] | Logit |
| | *Leaf number* | ii) Phenology [F] + Shade [F] + Phenology × Shade [F] + Upper AUDPC[F] + Lower AUDPC[F] + Herbivory[F] + Block [R] | Logit |

[1]$Log_{10}$-transformed

**Table S3.** The impact of phenology, shade, the interaction shade × phenology, the area under the disease progression curve (AUDPC) and leaf herbivory on the physiology, growth and survival of oak seedlings (*Quercus robur*).

| | Phenology | | | Shade | | | Shade × Phenology | | | Upper AUDPC | | | Lower AUDPC | | | Herbivory | | |
|---|---|---|---|---|---|---|---|---|---|---|---|---|---|---|---|---|---|---|
| Response variable | Df | F/$\chi^2$ | *P* | Df | F/$\chi^2$ | *P* | Df | F/$\chi^2$ | *P* | Df | F/$\chi^2$ | *P* | Df | F/$\chi^2$ | *P* | Df | F/$\chi^2$ | *P* |
| Leaf thickness | 2 | 1.8 | 0.41 | 1 | 5.45 | **0.021** | 2 | 0.43 | 0.8 | 1 | 0.025 | 0.87 | 1 | 2.41 | 0.12 | 1 | 7.4 | **0.006** |
| Chlorophyll | 2 | 32.15 | **<0.0001** | 1 | 7.41 | **0.006** | 2 | 3.31 | 0.19 | 1 | 2.67 | 0.1 | 1 | 0.22 | 0.64 | 1 | 0.78 | 0.19 |
| Seedling height | 2 | 53.19 | **<0.0001** | 1 | 0.66 | 0.42 | 2 | 1.63 | 0.44 | 1 | 3.77 | 0.052 | 1 | 1.97 | 0.16 | 1 | 0.83 | 0.36 |
| Leaf area | 2 | 56.75 | **<0.0001** | 1 | 0.57 | 0.45 | 2 | 0.23 | 0.89 | 1 | 1.91 | 0.17 | 1 | 1.91 | 0.17 | 1 | 1.2 | 0.27 |
| Leaf number | 2 | 38.42 | **<0.0001** | 1 | 0.001 | 0.97 | 2 | 0.99 | 0.61 | 1 | 0.44 | 0.51 | 1 | 1.86 | 0.17 | 1 | 0.43 | 0.51 |
| Survival | 2 | 35.19 | **<0.0001** | 1 | 0.55 | 0.45 | 2 | 0.07 | 0.96 | *X* | *X* | *X* | *X* | *X* | *X* | *X* | *X* | *X* |

**Table S4**. Repeated measures models of the proportion of leaves infected with powdery mildew and the severity of the infection on the upper and lower surface of *Quercus robur* seedlings leaves, as well as leaf herbivory, as a function of phenology, shade and date, and the interactions between phenology × shade, date × phenology, date × shade and date × phenology × shade. From weekly measurements during the period from 26 June 2018 to 31 July 2018.

| Upper surface powdery mildew proportion | Df | $\chi^2$ | P |
|---|---|---|---|
| Phenology | 2 | 6.89 | **0.032** |
| Shade | 1 | 1.24 | 0.265 |
| Date | 3 | 205.39 | **<0.0001** |
| Phenology × Shade | 2 | 2.48 | 0.29 |
| Date × Phenology | 6 | 97.85 | **<0.0001** |
| Date × Shade | 3 | 23.09 | **<0.0001** |
| Date × Phenology × Shade | 6 | 12.28 | 0.056 |

| Lower surface powdery mildew proportion | Df | $\chi^2$ | P |
|---|---|---|---|
| Phenology | 2 | 80.88 | **<0.0001** |
| Shade | 1 | 0.008 | 0.93 |
| Date | 3 | 306.71 | **<0.0001** |
| Phenology × Shade | 2 | 1.91 | 0.39 |
| Date × Phenology | 5 | 45.37 | **<0.0001** |
| Date × Shade | 3 | 16.63 | **<0.001** |
| Date × Phenology × Shade | 6 | 4.49 | 0.61 |

| Herbivory proportion | Df | $\chi^2$ | P |
|---|---|---|---|
| Phenology | 2 | 8.07 | **0.018** |
| Shade | 1 | 1.98 | 0.16 |
| Date | 1 | 1.93 | 0.17 |
| Phenology × Shade | 2 | 5.47 | 0.07 |
| Date × Phenology | 2 | 97.28 | **<0.0001** |
| Date × Shade | 1 | 0.31 | 0.58 |
| Date × Phenology × Shade | 2 | 5.23 | 0.073 |

| Upper surface powdery mildew severity | Df | $\chi^2$ | P |
|---|---|---|---|
| Phenology | 2 | 5.7 | 0.058 |
| Shade net | 1 | 1.47 | 0.23 |
| Date | 3 | 687.1 | **<0.0001** |
| Phenology × Shade net | 2 | 1.62 | 0.44 |
| Date × Phenology | 5 | 7.6 | 0.18 |
| Date × Shade | 3 | 12.22 | **0.007** |
| Date × Phenology × Shade | 5 | 2.92 | 0.71 |

| Lower surface powdery mildew severity | Df | $\chi^2$ | P |
|---|---|---|---|
| Phenology | 2 | 38.67 | **<0.0001** |
| Shade | 1 | 1.13 | 0.28 |
| Date | 2 | 87.81 | **<0.0001** |
| Phenology × Shade | 2 | 4.84 | 0.089 |
| Date × Phenology | 4 | 4.76 | 0.31 |
| Date × Shade | 2 | 2.76 | 0.25 |
| Date × Phenology × Shade | 6 | 0.34 | 0.84 |

| Herbivory severity | Df | $\chi^2$ | P |
|---|---|---|---|
| Phenology | 2 | 8.5 | **0.014** |
| Shade | 1 | 0.66 | 0.42 |
| Date | 1 | 0.21 | 0.65 |
| Phenology × Shade | 2 | 2.18 | 0.34 |
| Date × Phenology | 2 | 0.79 | 0.67 |
| Date × Shade | 1 | 0.21 | 0.65 |
| Date × Phenology × Shade | 2 | 0.95 | 0.62 |

**Table S5.** The impact of phenology, shade and their interaction on the severity of powdery mildew infection on the upper surface of the leaf, severity of powdery mildew infection on the lower surface of the leaf and leaf herbivory severity, on *Quercus robur* seedlings

| Response variable | Week Number | Phenology | | | Shade | | | Shade × Phenology | | |
|---|---|---|---|---|---|---|---|---|---|---|
| | | DF | F/$\chi^2$ | P | DF | F/$\chi^2$ | P | DF | F/$\chi^2$ | P |
| Upper leaf surface powdery mildew severity | Week 1 | 1 | 2.18 | 0.14 | 1 | 0.45 | 0.5 | 1 | 1.18 | 0.28 |
| | Week 2 | 2 | 5.63 | 0.059 | 1 | 1.96 | 0.16 | 2 | 0.56 | 0.75 |
| | Week 4 | 2 | 1.88 | 0.39 | 1 | 2.12 | 0.15 | 2 | 0.72 | 0.65 |
| | Week 6 | 2 | 0.66 | 0.72 | 1 | 8.97 | **0.0027** | 2 | 0.12 | 0.94 |
| Lower leaf surface powdery mildew severity | Week 2 | 2 | 5.6 | 0.06 | 1 | 1.96 | 0.16 | 2 | 0.56 | 0.75 |
| | Week 4 | 2 | 20.16 | **<0.0001** | 1 | 1.54 | 0.21 | 2 | 0.005 | 0.99 |
| | Week 6 | 2 | 3.33 | 0.19 | 1 | 0.26 | 0.61 | 2 | 2.67 | 0.26 |
| Leaf herbivory severity | Week 1 | 2 | 1.43 | 0.49 | 1 | 1.37 | 0.24 | 1 | 0.2 | 0.65 |
| | Week 4 | 2 | 5.29 | 0.071 | 1 | 0.17 | 0.68 | 2 | 1.85 | 0.4 |
